# SARS-CoV-2 Omicron BA.2.86: less neutralization evasion compared to XBB sub-variants

**DOI:** 10.1101/2023.09.26.559580

**Authors:** Yaling An, Xuemei Zhou, Lifeng Tao, Haitang Xie, Dedong Li, Ruyue Wang, Hua Hu, Zepeng Xu, Lianpan Dai, Kun Xu, George F. Gao

## Abstract

The continual emergence and circulation of new severe acute respiratory syndrome coronavirus 2 (SARS-CoV-2) variants have caused a great challenge for the coronavirus disease 2019 (COVID-19) pandemic control. Recently, Omicron BA.2.86 was identified with more than 30 amino acid changes on the spike (S) protein, compared to Omicron BA.2 or XBB.1.5. The immune evasion potential of BA.2.86 is of great concern. In this study, we evaluated the neutralizing activities of sera collected from participants and mice. Participants were divided into five groups according to their vaccination (inactivated vaccine, protein subunit vaccine ZF2001 or ZF2202-A) and infection (Omicron BF.7/BA.5.2) status. ZF2202-A is ZF2001 vaccine’s next-generation COVID-19 vaccine with updated bivalent Delta-BA.5 RBD-heterodimer immunogen. BALB/c mice were immunized with XBB.1.5 RBD-homodimer, BA.5-BA.2, Delta-XBB.1.5 or BQ.1.1-XBB.1.5 RBD-heterodimers protein vaccine candidates for evaluating the neutralizing responses. We found that Omicron BA.2.86 shows stronger immune evasion than BA.2 due to >30 additional mutations on S protein. Compared to XBB sub-variants, BA.2.86 does not display more resistance to the neutralizing responses induced by ZF2001-vaccination, BF.7/BA.5.2 breakthrough infection or a booster dose of ZF2202-A-vaccination. In addition, the mouse experiment results showed that BQ.1.1-XBB.1.5 RBD-heterodimer and XBB.1.5 RBD-homodimer induced high neutralizing responses against XBB sub-variants and BA.2.86, indicating that next-generation COVID-19 vaccine should be developed to enhance the protection efficacy against the circulating strains in the future.

## Introduction

Waves of severe acute respiratory syndrome coronavirus 2 (SARS-CoV-2) Omicron sub-variants circulate globally, including BA.1, BA.2, BA.5, BQ.1, XBB and their sub-variants (Fig. 1A). Recently, Omicron BA.2.86 was identified with more than 30 amino acid changes on the spike (S) protein, compared to BA.2 or XBB.1.5^1, 2^ (Fig. 1). The immune evasion potential of BA.2.86 is highly concerned. In addition, reported results from the preprint show controversial findings on whether BA.2.86 displays more resistance to the immune responses induced by breakthrough infection in the real world than XBB sub-variants^3, 4^. Therefore, we evaluated the neutralizing activities of sera collected from vaccination and infection for protection assessment.

**Fig. 1.**
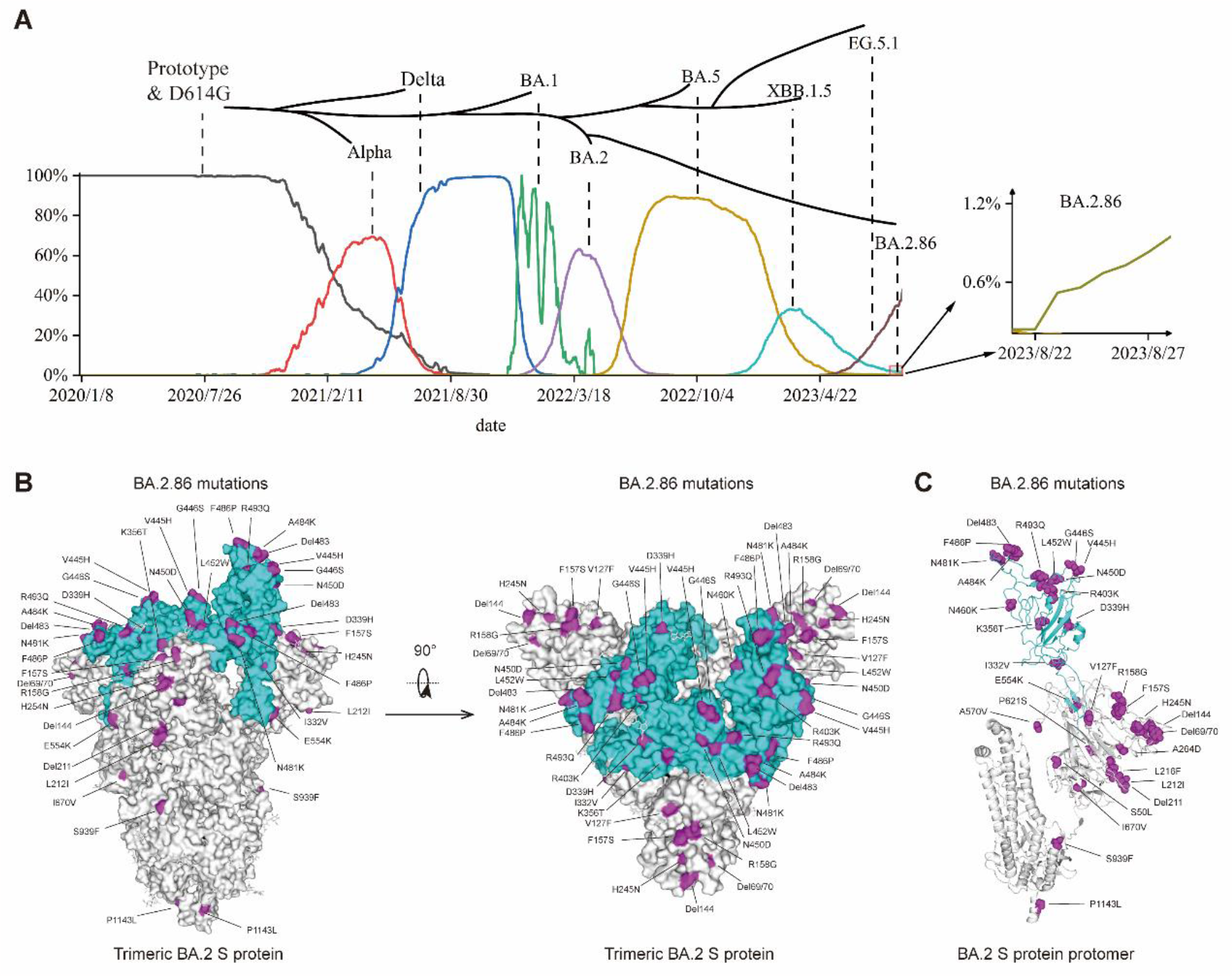
Waves of epidemic SARS-CoV-2 strains and the amino acid changes of BA.2.86 S protein. (A) The temporal evolution of the main epidemic variants and the emergence of BA. 2.86. The horizontal axis represents the timeline, the vertical axis represents the proportion of sequences, and the trend of each variant is represented by different colored lines. The branching diagram above represents the evolutionary relationship of each strain. Data were collected from PANGO lineage and GISAID. (B, C) The amino acid changes of BA.2.86 S protein compared to BA.2. BA.2 S protein is shown as trimer (B) and protomer (C) with gray color (PDB: 7XIW). RBD is shown as cyan color. Residues that changed in BA.2.86 are colored in purple.

In this study, the collected human sera samples were divided into five groups: (1) three homologous doses of COVID-19 inactivated vaccines (CoronaVac and BBIBP-CorV)^5, 6^; (2) three homologous doses of protein subunit vaccine ZF2001 (immunogen of SARS-CoV-2 RBD-homodimer)^7, 8^; (3-4) three doses of inactivated vaccines or ZF2001 vaccine followed by breakthrough infection during a wave of BF.7/BA.5.2 circulation in Beijing in December 2022^9^; (5) three doses of inactivated vaccines plus a booster of ZF2202-A vaccine. ZF2202-A is ZF2001 vaccine’s next-generation COVID-19 vaccine with updated bivalent Delta-BA.5 RBD-heterodimer immunogen^10^ and under clinical trial study (NCT05850507). Detailed information about the participants is available in Table 1. We measured the neutralization activities against SARS-CoV-2 using vesicular stomatitis virus backbone-based pseudotyped viruses^11^.

**Table 1.** Demographic and characteristics of participants.

| **Groups** | **Group 1** | **Group 2** | **Group 3** | **Group 4** | **Group 5** |
| --- | --- | --- | --- | --- | --- |
| **Status of vaccination or SARS-CoV-2 infections** | Three-dose inactivated vaccine | Three-dose ZF2001 | Three-dose inactivated vaccine plus breakthrough infection* | Three-dose ZF2001 plus breakthrough infection | Three-dose inactivated vaccine plus ZF2202-A booster |
| **No. of participants** | 24 | 8 | 25 | 25 | 12 |
| **Age, Years** |  |  |  |  |  |
| **Mean (SD)** | 32.0 (8.6) | 36.8 (9.0) | 29.7 (8.5) | 35.8 (14.4) | 53.8 (18.5), |
| **Median (IQR)** | 30.0 (25.0-36.8) | 35.5 (30.5-43.3) | 27.0 (24.0-30.0) | 31.0 (25.5-37.5) | 61.5 (36.3-68.0) |
| **Male, n (%)** | 13.0 (54.2%) | 1.0 (22.5%) | 6.0 (24.0%) | 11.0 (44.0%) | 6.0 (50.0%) |
| **Female, n (%)** | 11.0 (45.8%) | 7.0 (77.5%) | 19.0 (76.0%) | 14.0 (56.0%) | 6.0 (50.0%) |

\*Breakthrough infection were confirmed by antigen test for their nasopharyngeal swabs

## Results

Three homologous doses of inactivated vaccine-induced immune responses neutralize SARS-CoV-2 prototype and BA.2 pseudoviruses with geometric mean titers (GMTs) of 221 and 32, respectively, and lost neutralization activities against XBB sub-variants XBB.1.5/XBB.2.3/EG.5.1 and BA.2.86 with GMTs below the limit of detection (LOD) (Fig. 2A). The neutralizing GMTs induced by three homologous doses of ZF2001 vaccine were 2460 and 273 against prototype and BA.2 pseudoviruses, respectively (Fig. 2B). Compared to BA.2, BA.2.86 displayed increased immune evasion by a neutralization reduction factor of 8.0. The neutralization GMT against BA.2.86 was 34, higher than the GMTs against XBB.1.5/XBB.2.3/EG.5.1 (between 7 and 14) (Fig. 2B). BF.7/BA.5.2 breakthrough infection after inactivated or ZF2001 vaccine immunization elicited cross-reactive neutralizing antibody against BA.2.86 with GMTs of 32 and 197, respectively (Fig. 2C and 2D).

**Fig. 2.**
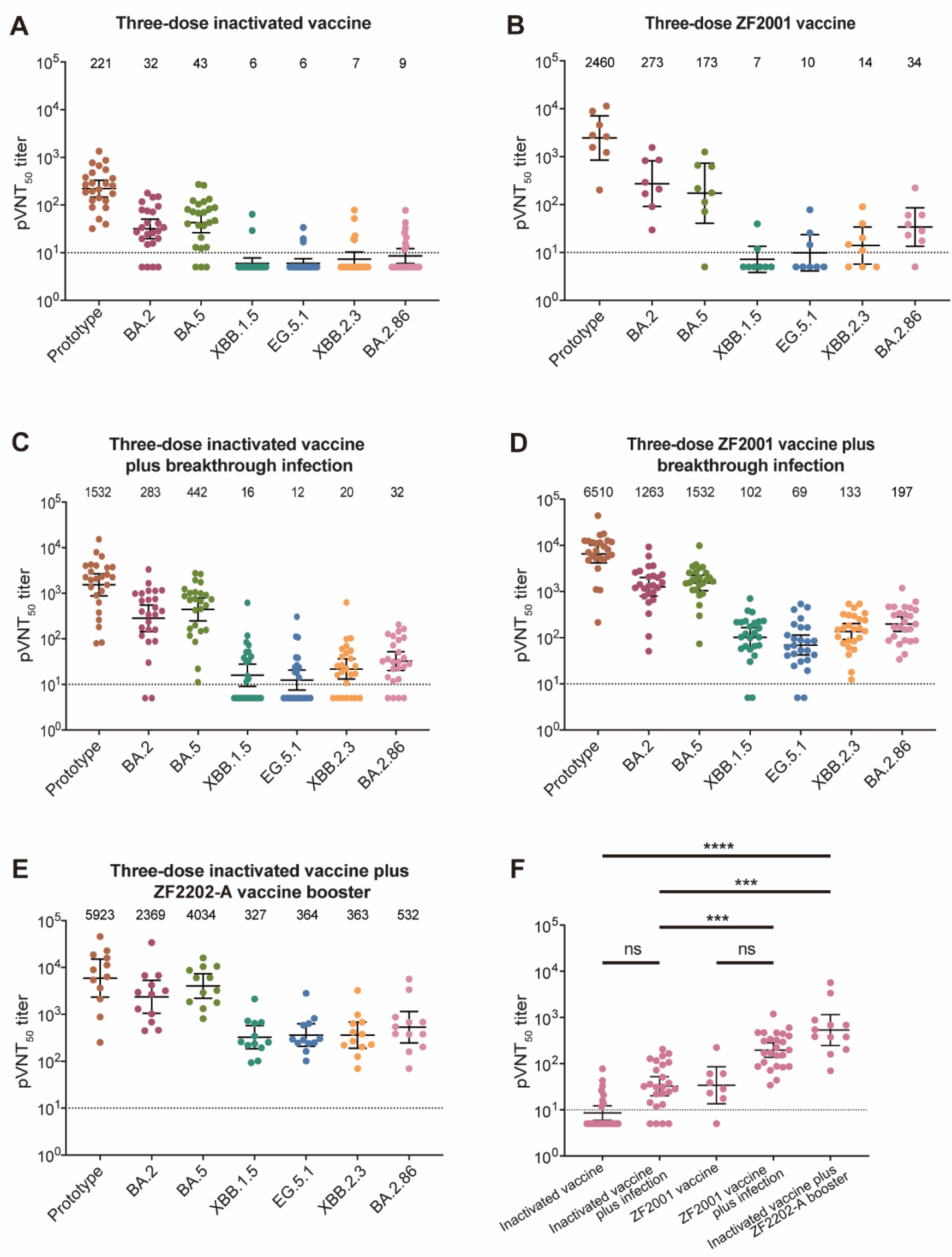
Neutralizing antibodies against SARS-CoV-2 variants by sera from humans. (A) The pVNT_50_ of sera from participants who had received three doses of inactivated vaccine. (B) The pVNT_50_ of sera from participants who had received three doses of ZF2001 vaccine. (C) The pVNT_50_ of sera from participants who had breakthrough infections during the late-2022 Omicron sub-variants BF.7 and BA.5.2 wave after three doses of inactivated vaccine. (D) The pVNT_50_ of sera from participants who had breakthrough infections during the late-2022 Omicron sub-variants BF.7 and BA.5.2 wave after three doses of ZF2001 vaccine. (E) The pVNT_50_ of sera from participants who had received three doses of inactivated vaccine and a booster of ZF2202-A vaccine. (F) p values were analyzed with a Kruskal-Wallis test first, and then Dunn’s multiple comparison test when the Kruskal-Wallis test was rejected. GMTs are shown at the top of each panel. The error bars indicate 95% confidence intervals. The dashed horizontal line indicates the LOD. pVNT_50_ below the LOD was determined as half the LOD.

A booster dose of the ZF2202-A vaccine after three doses of inactivated vaccine induced broad immune responses with 100% neutralization seropositive rates against all the tested pseudoviruses (Fig. 2E). The GMTs were 5923, 2369, 4034, 327, 364, 363 and 532 against prototype, BA.2, BA.5, XBB.1.5, EG.5.1, XBB.2.3 and BA.2.86, respectively (Fig. 2E). The neutralization activities against BA.2.86 induced by ZF2202-A vaccine boosting were higher than BF.7/BA.5.2 infection, indicating the potential protection efficacy of updated RBD-heterodimer vaccine against circulating SARS-CoV-2 strains (Fig. 2F).

Next, we evaluated the neutralizing profiles induced by several other vaccine candidates in a mouse model, including XBB.1.5 RBD-homodimer, BA.5-BA.2, Delta-XBB.1.5 and BQ.1.1-XBB.1.5 RBD-heterodimers (Fig. 3). Prototype RBD-homodimer and Delta-BA.5 RBD-heterodimer were immunized as a comparison (Fig. 3). As expected, XBB.1.5 RBD-homodimer, Delta-XBB.1.5 and BQ.1.1-XBB.1.5 RBD-heterodimers elicited high levels of neutralizing antibodies against XBB sub-variants, with GMTs of ∼10^4^ (Fig. 3D-3F). For neutralizing BA.2.86, XBB.1.5 RBD-homodimer and BQ.1.1-XBB.1.5 RBD-heterodimer induced potent cross-reactivity with GMTs of 3820 and 4801, respectively (Fig. 3E and 3F). The neutralizing antibodies induced by BA.5-BA.2 RBD-heterodimer were high against Omicron BA.2, BA.5, BF.7 and BQ.1.1 with GMTs between 37183 and 59428, which decreased to a range between 503 and 1199 against XBB sub-variants and BA.2.86 (Fig. 3C). The results from mouse experiments suggested that the next-generation COVID-19 vaccine should be developed with updated SARS-CoV-2 sequences to confront the surge of newly emerged variants.

**Fig. 3.**
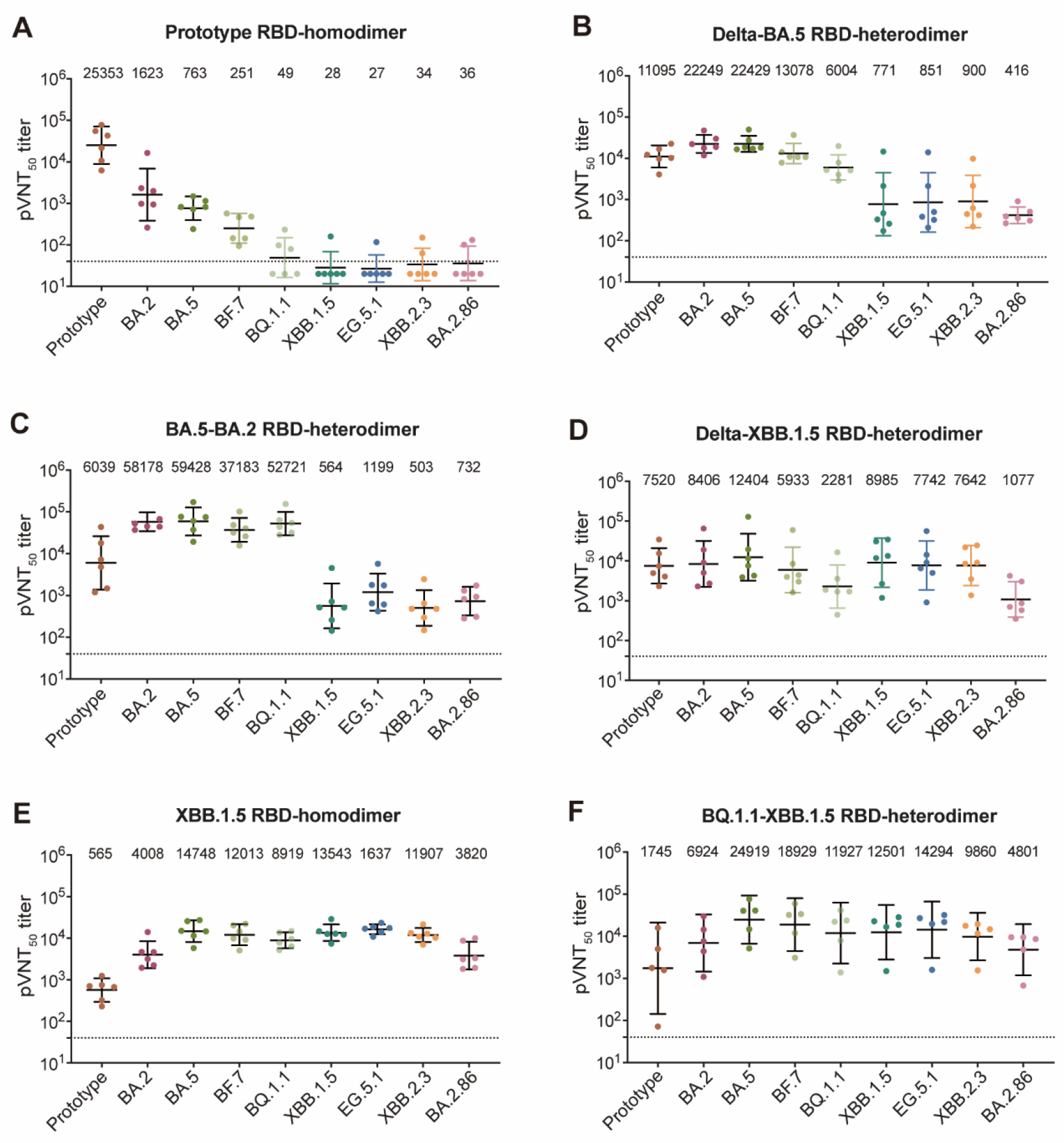
Neutralizing antibodies against SARS-CoV-2 variants by sera from mice. BALB/c mice were immunized with three doses of prototype RBD-homodimer (A), Delta-BA.5 RBD-heterodimer (B), BA.5-BA.2 RBD-heterodimer (C), Delta-XBB.1.5 RBD-heterodimer (D), XBB.1.5 RBD-homodimer (E) or BQ.1.1-XBB.1.5 RBD-heterodimer (F) using AddaVax as adjuvant. PBS plus adjuvant was given as the sham control. Sera were collected at 14 days post the third immunization. The pVNT_50_ of murine sera were measured. The details of mice experiments were described in the Materials and Methods section. GMTs are shown at the top of each panel. The error bars indicate 95% confidence intervals. The dashed horizontal line indicates the LOD. pVNT_50_ below the LOD was determined as half the LOD.

## Conclusion and Discussion

In conclusion, this study reveals that Omicron BA.2.86 shows stronger immune evasion than BA.2 due to extra mutations. In the real world, the ZF2001 vaccine-immunization and BF.7/BA.5.2-breakthrough infection induced higher sera neutralizing levels against BA.2.86 than the recently-circulating XBB sub-variants strains XBB.1.5, XBB.2.3 and EG.5.1^12^, suggesting that BA.2.86 does not harbor more resistance to the herd immunity induced by immunization and breakthrough infection. In addition, the COVID-19 vaccine ZF2202-A (delta-BA.5 RBD-heterodimer) boosts neutralizing responses against Omicron sub-variants in humans, including EG.5.1 and BA.2.86. Next-generation vaccine candidates, such as BQ.1.1-XBB.1.5 RBD-heterodimer and XBB.1.5 RBD-homodimer, elicit high neutralizing responses against XBB sub-variants and BA.2.86 and deserve further clinical development.

COVID-19 protein subunit vaccine ZF2001, based on SARS-CoV-2 prototype RBD-homodimer, is demonstrated to be safe, immunogenic and protective^8, 13^. Due to immune evasion of SARS-CoV-2 variants, RBD-homodimer is engineered by combining RBDs from two strains as bivalent vaccine to induce broad immune responses^10^. ZF2001’s second-generation vaccine ZF2202 contains Delta-BA.1 RBD-heterodimer immunogen and induces potent antibody responses in humans^14^. However, SARS-CoV-2 continuously evolves to escape the herd immunity. ZF2202 is further updated with the immunogen of Delta-BA.5 RBD-heterodimer (ZF2202-A) to enhance the protection efficacy against the circulating SARS-CoV-2 strains. In the future, the COVID-19 vaccine can be annually inoculated like the current influenza vaccine. A combination of the updated COVID-19 and influenza vaccines will reduce the inoculation frequency and protect against these two respiratory viruses, which is worth exploring^15-17^.

## Materials and Methods

### Human serum samples

Human serum samples were classified into five groups in this study. The first group participants received three doses of inactivated vaccine (CoronaVac or BBIBP-CorV)^5, 6^. The second group received three doses of recombinant protein subunit vaccine (ZF2001)^7,8^. The third group participants received three doses of inactivated vaccine and had breakthrough infection during the BF.7/BA.5.2 wave in late 2022 in Beijing, China^9^. The fourth group participants received three doses of ZF2001 vaccine and had breakthrough infection during the BF.7/BA.5.2 wave in late 2022 in Beijing, China. The fifth group received three doses of inactivated vaccine, followed by boosting with one dose of Delta-Omicron-BA.5 RBD-heterodimer protein vaccine (ZF2202-A). Serum samples from the fifth group were from a clinical trial (NCT05850507), and were approved by the clinical research ethics board of the First Affiliated Hospital of Wannan Medical College (Yijishan Hospital) ([2023]KY34). Serum samples from the first group were provided by Anhui Zhifei Longcom Biopharmaceutical. Serum samples from the second, third and fourth groups were collected from the participants in real-world. All participants signed the written informed consent. Detailed information is available in Table 1.

### Pseudotyped virus neutralization assay

The pseudotyped virus displaying SARS-CoV-2 (prototype and variants) S protein expresses GFP in infected cells. They were prepared as previously described^18^. Serially diluted mice sera were incubated with pseudotyped virus at 37°C for 1 hour, then the sera-virus mixtures were transferred to pre-plated Vero E6 cell monolayers in 96-well plates. After incubation for 15 hours, the transducing unit numbers were calculated on a CQ1 confocal image cytometer (Yokogawa). Neutralization titer was determined by fitting nonlinear regression curves using GraphPad Prism and calculating the reciprocal of the serum dilution required for 50% neutralization of infection. Neutralization titer below the limit of detection was determined as half the limit of detection.

### Mouse experiments

Specific pathogen-free (SPF) female BALB/c mice were purchased from Beijing Vital River Laboratory Animal Technology Co., Ltd. (licensed by Charles River). All mice were allowed free access to water and standard chow diet and provided with a 12-hour light and dark cycle (temperature: 20-25°C, humidity: 40%-70%). All mice used in this study are in good health and are not involved in other experimental procedure. They were housed under SPF conditions in the laboratory animal facilities at IMCAS. The mice experiments conducted in IMCAS were approved by the Committee on the Ethics of Animal Experiments of the IMCAS, and performed in compliance with the recommendations in the Guide for the Care and Use of Laboratory Animals of the IMCAS Ethics Committee.

Antigen proteins were prepared as previously described^10^. AddaVax adjuvant was purchased from InvivoGen. Groups of 6-to 8-week-old female BALB/c mice were immunized with three doses of 2-μg immunogens adjuvanted by AddaVax. The interval between the first and second doses was 21 days. The interval between the second and third doses was 21 days. The serum samples were collected 14 days after the last immunization.

### Data and statistical analyses

The numbers of pseudovirus-infected cells were obtained using a CQ1 confocal image cytometer. Each sample was tested in duplicate. The pVNT_50_ was determined by fitting non-linear regression curves (log(inhibitor) vs. normalized response -Variable slope) using GraphPad Prism (8.0.1) and calculating the reciprocal of the serum dilution required for 50% neutralization of infection. The model of the non-linear regression curve is Y=100/(1+10^((LogpVNT50-X)*HillSlope))). HillSlope describes the steepness of the family of curves. The neutralizing antibody titers were illustrated via GMT with 95% CI (geometric mean titer with 95% confidence interval). p values were analyzed with Kruskal-Wallis test first, and then Dunn’s multiple comparison test when the Kruskal-Wallis test was rejected.

## Acknowledgments

This work is supported by the National Key R&D Program of China (2020YFA097100 and 2021YFC2302600) and a grant from the Bill & Melinda Gates Foundation (INV-027420). L.D. is supported by the Excellent Young Scientist Program from National Natural Science Foundation of China (NSFC) (82122031) and Youth Innovation Promotion Association CAS, China (2018113). K.X. is supported by the NSFC (82202030) and Young Elite Scientists Sponsorship Program by CAST (2022QNRC001). We thank all volunteers for providing blood samples and Anhui Zhifei Longcom Biopharmaceutical for their help in providing sera samples. We thank Jialin Fan (University of Chinese Academy of Sciences) for his assistance in the experiments.

## Author Contributions

L.D., K.X. and G.F.G. conceived and coordinated the study. Y.A., X.Z. and D.L. conducted the experiments. H.X. and H.H. acquired serum samples for the clinical trial. L.T. and R.W. organized the investigational products. Y.A., X.Z., Z.X., L.D. and K.X. analyzed the data. K.X. drafted the manuscript. G.F.G. revised the manuscript.

## Competing Interests

Y.A., L.D, K.X. and G.F.G. are listed in the patent as the inventors of the prototype RBD-dimer as coronavirus vaccines. L.D., K.X. and G.F.G are listed in the patent as the inventors of Delta-Omicron RBD-dimer as coronavirus vaccine. L.T. and R.W. are employees at Anhui Zhifei Longcom Biopharmaceutical Co. Ltd. All other authors declare no competing interests.

## References

1 Rasmussen M, Møller FT, Gunalan V et al. First cases of SARS-CoV-2 BA.2.86 in Denmark, 2023. Euro Surveill 2023; 28.

2 Looi MK. Covid-19: Scientists sound alarm over new BA.2.86 “Pirola” variant. BMJ 2023; 382:1964.

3 Yang S, Yu Y, Jian F et al. Antigenicity and infectivity characterization of SARS-CoV-2 BA.2.86. bioRxiv 2023:DOI: 10.1101/2023.1109.1101.555815.

4 Qu P, Xu K, Faraone JN et al. Immune Evasion, Infectivity, and Fusogenicity of SARS-CoV-2 Omicron BA.2.86 and FLip Variants. bioRxiv 2023:DOI: 10.1101/2023.1109.1111.557206.

5 Jara A, Undurraga EA, Gonzalez C et al. Effectiveness of an inactivated SARS-CoV-2 vaccine in Chile. N Engl J Med 2021; 385:875–884.

6 Al Kaabi N, Zhang Y, Xia S et al. Effect of 2 inactivated SARS-CoV-2 vaccines on symptomatic COVID-19 infection in adults: a randomized clinical trial. JAMA 2021; 326:35–45.

7 Dai L, Zheng T, Xu K et al. A universal design of betacoronavirus vaccines against COVID-19, MERS, and SARS. Cell 2020; 182:722–733 e711.

8 Dai L, Gao L, Tao L et al. Efficacy and safety of the RBD-Dimer-based COVID-19 vaccine ZF2001 in adults. N Engl J Med 2022; 386:2097–2111.

9 Pan Y, Wang L, Feng Z et al. Characterisation of SARS-CoV-2 variants in Beijing during 2022: an epidemiological and phylogenetic analysis. Lancet 2023; 401:664–672.

10 Xu K, Gao P, Liu S et al. Protective prototype-Beta and Delta-Omicron chimeric RBD-dimer vaccines against SARS-CoV-2. Cell 2022; 185:2265–2278 e2214.

11 Nie J, Li Q, Wu J et al. Establishment and validation of a pseudovirus neutralization assay for SARS-CoV-2. Emerg Microbes Infect 2020; 9:680–686.

12 Abbasi J. What to Know About EG.5, the Latest SARS-CoV-2 “Variant of Interest”. JAMA 2023.

13 Yang S, Li Y, Dai L et al. Safety and immunogenicity of a recombinant tandem-repeat dimeric RBD-based protein subunit vaccine (ZF2001) against COVID-19 in adults: two randomised, double-blind, placebo-controlled, phase 1 and 2 trials. Lancet Infect Dis 2021; 21:1107–1119.

14 Dai L, Duan H, Liu X et al. Omicron neutralisation: RBD-dimer booster versus BF.7 and BA.5.2 breakthrough infection. Lancet 2023; 402:687–689.

15 Izikson R, Brune D, Bolduc JS et al. Safety and immunogenicity of a high-dose quadrivalent influenza vaccine administered concomitantly with a third dose of the mRNA-1273 SARS-CoV-2 vaccine in adults aged >/=65 years: a phase 2, randomised, open-label study. Lancet Respir Med 2022; 10:392–402.

16 Toback S, Galiza E, Cosgrove C et al. Safety, immunogenicity, and efficacy of a COVID-19 vaccine (NVX-CoV2373) co-administered with seasonal influenza vaccines: an exploratory substudy of a randomised, observer-blinded, placebo-controlled, phase 3 trial. The Lancet Respiratory Medicine 2021; 10:167–179.

17 Li Y, Liu P, Hao T et al. Rational design of an influenza-COVID-19 chimeric protective vaccine with S-RBD and HA-stalk. Emerg Microbes Infect 2023:2231573.

18 Zhao X, Li D, Ruan W et al. Effects of a prolonged booster interval on neutralization of Omicron variant. N Engl J Med 2022; 386:894–896.

